# Lateral Membrane Heterogeneity Shapes Antimicrobial Association and Convergent Mechanical Responses

**DOI:** 10.64898/2026.11.08.765177

**Authors:** Mateusz Rzycki, Naresh Yandrapalli, Dominik Drabik

**Affiliations:** Department of Biomedical Engineering, Wroclaw University of Science and Technology, Wroclaw, 50-370, Poland; Department of Synthetic Biology, University of Saarland, Saarbrücken, Germany

**Keywords:** antimicrobial agents, molecular dynamics, microfluidics, bacterial domains, octenidine, chlorhexidine, polymyxin B

## Abstract

Bacterial membranes are laterally heterogeneous, creating local physicochemical environments that can shape interactions with membrane-active antimicrobials. However, whether this spatial organization only affects local drug association or propagates into collective membrane responses remains unclear. Here we show that lateral membrane heterogeneity shapes antimicrobial association while distinct molecular responses converge on a common loss of membrane mechanical stability. Combining coarse-grained molecular dynamics, giant unilamellar vesicle fluctuation spectroscopy and microfluidic trapping, we follow octenidine, chlorhexidine and polymyxin B from molecular association to vesicle-scale behaviour. All three compounds preferentially associate with domain-associated membrane environments, but differ in how this preference changes with loading and in their effects on lipid mobility. Despite these differences, all three shift the membrane toward a mechanically softer state, while increasing exposure accelerates loss of vesicle stability under flow. These findings identify lateral membrane heterogeneity as an active determinant of antimicrobial response and provide a physical framework for understanding membrane-targeting activity beyond bulk composition and membrane-averaged charge.

## Introduction

The bacterial membrane is not a laterally uniform lipid matrix. In addition to providing a permeability barrier, it is organized into membrane regions that differ in lipid composition, packing, and possess physicochemical properties, contributing to processes including signal transduction, protein organization, transport, and adaptation to environmental stress.^1–3^ Such functional lateral organization can arise from interactions between membrane proteins and specific lipid species, but also from the intrinsic tendency of particular lipids to form compositionally distinct environments.^4^ Hopanoids are particularly relevant in this context. These bacterial polycyclic lipids share several membrane-ordering properties with sterols and can promote ordered lipid environments in model membranes.^5^ Consequently, molecules approaching a bacterial membrane encounter a spatially heterogeneous physicochemical landscape rather than a single uniform membrane phase.

Membrane heterogeneity may be particularly important for active antimicrobial compounds (see Fig.1A). Their interaction is commonly discussed in terms of electrostatic attraction, hydrophobic insertion, membrane permeabilization, or changes in lipid packing.^6–8^ However, all of these processes depend on local membrane composition and organization. Differences in surface charge, lipid order, packing density, and lateral mobility can change both the accessibility of the membrane and the energetic cost of antimicrobial association. Consistent with this view, coarse-grained simulations of phase-separated bilayers showed that several antimicrobial peptides preferentially partition into liquid-disordered regions, with some additionally accumulating at domain interfaces depending on lipid packing and peptide loading.^9^ Experimental SANS and SAXS measurements further showed that antimicrobial peptides can reorganize pre-existing domains and alter their line tension through phase-dependent changes in bilayer thickness.^10^ Similar coupling between antimicrobials binding and lateral membrane organization has been observed in bacterial-mimetic systems, where cationic molecules cluster anionic lipids or promote lipid segregation, ^8^ and solid-state NMR has revealed peptide-dependent differences in lipid ordering in PE/PG membranes under phase-coexistence conditions.^11^ Although these systems differ chemically from the compounds studied here, they point to the same general principle: lateral membrane organization can influence where membrane-active molecules accumulate, while their binding can in turn reshape the local lipid environment. A related response has also been observed in intact bacteria, where daptomycin reorganized fluid lipid domains in *B. subtilis* and displaced membrane-associated cell-envelope biosynthetic machinery.^12^ Therefore, membrane heterogeneity may not only influence the initial association of antimicrobial molecules but may also determine how their local interactions propagate into larger-scale changes in membrane dynamics and mechanics.

**Figure 1:**
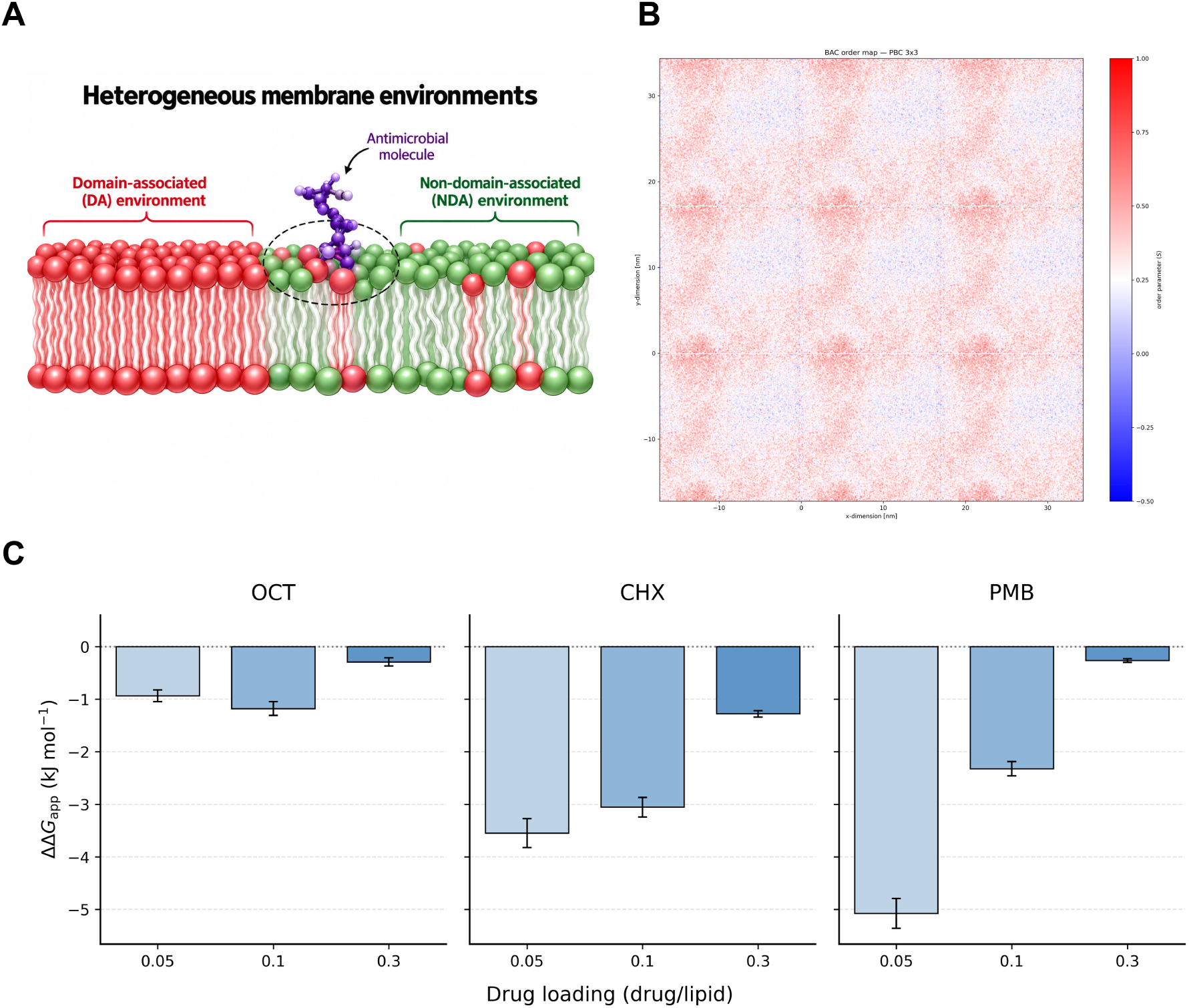
Lateral membrane organization shapes antimicrobial association. (A) Schematic representation of the domain-associated (DA; PC, PG, CL and HOP) and non-domain-associated (NDA; PE) membrane environments sampled by an antimicrobial molecule. (B) Representative *S*_CD_-derived order map of the untreated membrane, displayed with periodic replication to visualize the continuity of its lateral organization. (C) Relative apparent compositional preference free energy, ΔΔ*G*_app_, for OCT, CHX, and PmB at increasing drug-to-lipid ratios. Negative values indicate preferential association with the DA environment. Error bars represent temporal uncertainties estimated using block averaging.

This aspect remains insufficiently resolved because many experimental and computational studies of membrane-active antimicrobials employ compositionally homogeneous bilayers or interpret membrane responses primarily through spatially averaged properties. This is particularly relevant for compounds whose activity cannot be reduced to a simple electrostatic interaction with negatively charged lipids. For example, polymyxin B (PmB) exhibits strongly composition-dependent membrane behaviour. Its interaction is affected by both membrane charge and lipid packing, and simulations of different bacterial membrane models have reported changes in lipid diffusion, bending rigidity, lipid asymmetry, and lateral phase organization.^13–15^ Similarly, our previous studies of octenidine (OCT) and chlorhexidine (CHX) showed that their membrane activity cannot be explained by electrostatic selectivity alone.^6^ OCT was more strongly associated with changes in membrane mechanical properties, whereas CHX produced more discrete alterations of membrane organization. Importantly, the membrane models used in that study were homogeneous and therefore could not address how lateral heterogeneity contributes to these interactions. We therefore hypothesized that lateral membrane heterogeneity creates distinct physicochemical environments that differentially modulate antimicrobial association and thereby influence the subsequent membrane response.

To test this hypothesis, we investigated three structurally and functionally distinct cationic antimicrobials: OCT, CHX, and PmB using an *Methylorubrum extorquens*-like membrane model displaying spontaneous lateral heterogeneity. ^16–18^ Rather than assigning antimicrobial action to a single membrane property, we followed the response across complementary spatial scales. Coarse-grained molecular dynamics simulations were used to determine preferential association with distinct lateral lipid environments and the accompanying changes in lipid mobility and membrane mechanics. These observations were complemented by fluctuation spectroscopy of giant unilamellar vesicles, providing a measure of the collective mechanical state of the heterogeneous membrane, and by microfluidic trapping experiments probing the stability of whole vesicles under confinement.

Our results show that lateral membrane organization provides a common basis for antimicrobial association while the subsequent membrane response remains compound-dependent. All three compounds displayed preferential association with domain-associated lipid environments, although the strength and concentration dependence of this preference differed substantially. CHX maintained preferential association over the investigated dosing range and progressively restricted lipid mobility, whereas the initial preference of OCT weakened with increase in loading and was accompanied by increasingly pronounced membrane remodeling. The membrane-associated PmB population showed the strongest compositional preference and a distinct effect on membrane dynamics. Despite these different molecular-scale behaviours, all three compounds shifted the membrane toward lower bending rigidity, a response observed independently in simulations and GUV fluctuation measurements. These changes were further reflected at the level of the entire vesicle, as increasing antimicrobials exposure accelerated GUV loss from microfluidic trapping structures, indicating reduced mechanical stability of the treated membranes. Together, our results connect preferential interactions with lateral membrane organization to changes in lipid dynamics, collective membrane mechanics, and ultimately whole-vesicle stability. They therefore position lateral membrane heterogeneity as an active determinant of antimicrobial response and provide a framework for understanding membrane-targeting activity beyond homogenous lipid composition alone.

## Materials and Methods

### Lipids, antimicrobial agents, and reagents

Phospholipids were purchased from Avanti Polar Lipids and included POPC (1-palmitoyl-2-oleoyl-sn-glycero-3-phosphocholine), DPPG (1,2-dipalmitoyl-sn-glycero-3-phospho-(1’-racglycerol)), DiPE (1,2-dilinoleoyl-sn-glycero-3-phosphoethanolamine), and 18:1 cardiolipin (CL). The hopanoid 22-hydroxyhopane (HOP) was obtained from Chiron. Atto488-DOPE (Sigma-Aldrich) and DiIC_12_ (Thermo Fisher Scientific) were used for fluorescence visualization. All aqueous solutions were prepared using ultrapure water. Octenidine (OCT) and polymyxin B (PmB) were obtained from Thermo Fisher Scientific as powders and dissolved in water before use. Chlorhexidine digluconate (CHX) was purchased from Sigma-Aldrich as an aqueous solution and diluted to the required final concentrations. The concentration ranges were selected to span low, intermediate, and high antimicrobial exposure regimes informed by concentrations reported in antimicrobial susceptibility, membrane-interaction, and application-oriented studies.^19–24^ Rather than corresponding to a single organism-specific MIC, the selected ranges were intended to resolve concentration-dependent membrane responses across the three structurally distinct compounds.

### Giant unilamellar vesicles

Giant unilamellar vesicles (GUVs) were prepared using the *M. extorquens*-like membrane composition established and biophysically characterized in our previous work^18^ to reproduce key compositional features of the *M. extorquens* membrane.^16,17^ The membrane contained POPC, DiPE, DPPG, HOP, and CL at a molar ratio of 0.30:0.30:0.20:0.18:0.02, respectively. For fluorescence visualization, the lipid mixture was supplemented with 1 mol% Atto488-DOPE and 1 mol% DiIC_12_. Lipids dissolved in chloroform were mixed to a total lipid concentration of 1 mM, and 50 *µ*L of the mixture was distributed along platinum electrodes. Following solvent evaporation, residual chloroform was removed under vacuum for 3 h. The electrodes were subsequently mounted in a glass electroformation chamber maintained at 45 *^◦^*C and immersed in 120 mOsm sucrose solution. GUV formation was induced using a 10 Hz square-wave alternating electric field. The voltage was initially set to 1 V and increased by 1 V h*^−^*^1^ until reaching 4 V, after which electroformation was continued for a total of 24 h. For membrane mechanical measurements, GUVs were exposed to OCT at 20, 200, and 600 *µ*M, CHX at 20, 50, and 200 *µ*M, and PmB at 0.5, 1, 2, 20, and 50 *µ*M. Untreated vesicles of identical lipid composition were used as reference samples.

### Fluorescence microscopy and vesicle fluctuation spectroscopy

Membrane bending rigidity was determined from thermally induced shape fluctuations of individual GUVs within the Helfrich elastic membrane framework. Time-resolved fluorescence images were acquired using a Stellaris confocal microscope (Leica, Wetzlar, Germany) equipped with an HC PL APO 86*×*/1.20 water-immersion objective. Membranes were labeled with Atto488-DOPE and DiIC12, and fluorescence excitation was provided by the white-light laser, using excitation wavelengths of 488 nm for Atto488-DOPE and 549 nm for DiIC12. Image sequences were recorded at 256 *×* 256 pixels using a hybrid detector (HyD; Leica), with pixel sizes ranging from 0.06 to 0.12 *µ*m, depending on the applied optical zoom. The corresponding frame integration times ranged from 130 to 388 ms. Fluorescence excitation was provided by the white-light laser set to 488 nm. Individual fluctuation recordings typically consisted of 1000 consecutive images.

The imaging plane was positioned close to the equatorial section of each vesicle to maximize contour fidelity. For every recorded frame, the membrane contour was identified, and the instantaneous radial displacement was extracted as a function of polar angle. Contour detection was performed using custom-written image-analysis software and applied consistently to all experimental conditions. The resulting time-dependent contour fluctuations were subsequently analysed using the statistical approach described previously.^25^ Measurements were initiated approximately 15 min after antimicrobial addition, and at least three independent experimental repeats were performed for each antimicrobial agent.

### Microfluidic device operation

The fabrication and geometry of the microfluidic trapping device have been described in detail previously.^26^ Before GUV loading, the microchannels were flushed with 2% Pluronic F127 solution via centrifugation and incubated for 30 min at room temperature to minimize nonspecific vesicle adhesion to the channel walls and trapping structures. The GUV suspension was subsequently introduced into the device, followed by centrifugation for 10 min to facilitate vesicle preloading into the traps. The system was then operated at a flow rate of 5 *µ*L min*^−^*^1^, and approximately 300–500 *µ*L of GUV solution was passed through the device to establish stable flow conditions and occupy the traps. Before introducing the drug, the flow was briefly stopped, and the reservoir was rinsed twice with fresh solution. Antimicrobial exposure was initiated by perfusing approximately 180 *µ*L of the corresponding drug solution through the chip at the same flow rate. All solutions used during the experiments were osmotically matched to minimize vesicle deformation or rupture arising from osmotic imbalance.

Microfluidic trapping experiments were imaged using a Leica TCS SPE microscope (Leica, Wetzlar, Germany) equipped with a 63*×*/1.30 ACS APO oil-immersion objective. Fluorescence was excited using a 488 nm laser. In contrast to the Stellaris-based measurements described above, the microfluidic experiments were acquired using this single excitation wavelength and were intended primarily for visualization and tracking of the GUV membrane during trapping experiments rather than for spectrally resolved imaging of the two membrane probes. At least two independent experimental repeats were performed for each antimicrobial agent.

### Coarse-grained molecular dynamics simulations

Coarse-grained molecular dynamics simulations were based on the *M. extorquens*-like membrane model previously developed and characterized in detail.^16–18^ The bilayer contained 1000 lipids with the same composition as the experimental system: 30% POPC, 30% DiPE, 20% DPPG, 18% HOP, and 2% CL. The membrane systems had been equilibrated previously in the absence of antimicrobial agents and were used here as the starting configurations for the drug-treated simulations. OCT, CHX, and PmB coarse-grained representations were prepared on the basis of previously published molecular topologies and adapted to the forcefield representation employed in the present simulations.^14,27^ OCT and CHX each carried a net charge of +2*e*, whereas PmB carried a net charge of +5*e*. For each compound, antimicrobial molecules were initially distributed in the aqueous phase on both sides of the pre-equilibrated membrane using custom Python scripts. The number of molecules was adjusted to obtain antimicrobial-to-lipid ratios of 0.05, 0.10, and 0.30, corresponding to 50, 100, and 300 antimicrobial molecules per 1000-lipid bilayer, respectively. Systems contained approximately 0.05 M NaCl, and additional counterions were introduced to neutralize the antimicrobial charges and maintain overall system electroneutrality. The systems contained, on average, approximately 22 polarizable Martini water beads per lipid, corresponding to roughly 90 water molecules per lipid in an atomistic representation.

Following antimicrobial placement, all systems underwent the same equilibration procedure as the corresponding drug-free membrane described previously. ^18^ Production simulations were then performed for approximately 5 *µ*s. Unless otherwise stated, quantitative analyses were conducted over the final 500 ns of each trajectory, for which the area per lipid and membrane thickness converged across all investigated systems (Fig. S1). Simulations were performed using GROMACS 2022.4^28^ with Martini 2.3P force field^29^ and polarizable water. A 20 fs integration time step was employed. The systems were simulated in the NPT ensemble at 295 K using the V-rescale thermostat with a coupling constant of 1 ps. Semi-isotropic pressure coupling was applied using the Parrinello–Rahman barostat, ^30^ with a coupling constant of 12 ps and a compressibility of 3 *×* 10*^−^*^4^ bar*^−^*^1^.

Membrane structural, dynamic, and mechanical properties, including lateral diffusion, area per lipid, membrane thickness, lipid order, interdigitation and bending rigidity, were determined using the analysis framework established previously.^18^ Trajectory processing and additional analyses were performed using custom Python scripts based on the MDAnalysis library.^31^

### Local lipid enrichment and apparent compositional preference free energy

Preferential antimicrobial association was quantified from the local lipid composition surrounding individual drug molecules. Each antimicrobial molecule was analysed independently and assigned to the nearest membrane leaflet. Lipids located within a lateral distance of 0.6 nm from the antimicrobial molecule were used to define its local membrane environment. For each lipid species, the local mole fraction (*x^local^*, the fraction of lipid *i* within the defined local neighbourhood) was normalized to its abundance in the corresponding leaflet (*x^ref^* , the leaflet-wide mole fraction of lipid *i*), yielding a compositional enrichment factor:

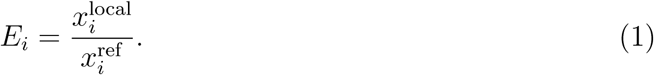

The classification of lateral membrane environments was based on the membrane organization established independently in our previous characterization of the untreated system.^18^ PC, PG, CL and HOP were treated as the domain-associated lipid pool, whereas PE formed the non-domain-associated lipid pool. This classification therefore represents a composition-based descriptor of the previously characterized membrane lateral organization rather than a fixed assignment of individual lipid molecules to a thermodynamic phase (see Fig. 1B). The enrichment factors were transformed into an apparent compositional preference free energy:

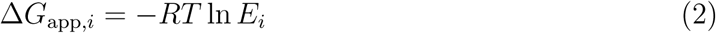

where *R* is the gas constant and *T* is the simulation temperature. Negative Δ*G*_app_*_,i_* values therefore correspond to local enrichment of lipid *i*, whereas positive values indicate depletion. Thus the relative preference between the two membrane environments was expressed as:

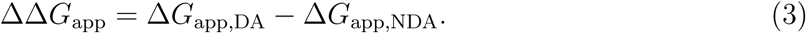

where Δ*G*_app,DA_ and Δ*G*_app,NDA_ describe the apparent preference for the domain-associated and non-domain-associated lipid pools, respectively. Negative values of ΔΔ*G*_app_ indicate preferential association with the domain-associated environment, and positive values indicate a relative preference for the non-domain-associated environment. Because this quantity is derived from compositional enrichment rather than from a direct equilibrium free-energy calculation, it is reported as an apparent compositional preference free energy rather than an absolute binding free energy. Local lipid fractions were first averaged across individual antimicrobial molecules within each trajectory frame and subsequently averaged over the analysed trajectory interval. Temporal uncertainties were estimated using block averaging.

## Results and Discussion

### Antimicrobials exhibit distinct preferences for lateral membrane environments

We hypothesized that lateral membrane heterogeneity creates distinct physicochemical environments that differentially modulate antimicrobial association and thereby influence the subsequent membrane response. We therefore first examined whether OCT, CHX, and PmB interact uniformly with the heterogeneous bilayer or preferentially associate with particular lateral lipid environments. The DA/NDA classification was based on the lateral ordering previously resolved for the untreated membrane using lipid-specific *S*_CD_ maps.^18^ Regions associated with PC, PG, CL and HOP accounted for the ordered, domain-associated environment, whereas PE represented the remaining non-domain-associated lipid pool. The apparent compositional preference free energy (ΔΔ*G*_app_) was used to compare antimicrobial preference between these environments, with negative values indicating preferential association with the domain-associated environment (Fig. 1)A,C.)

All three antimicrobial agents displayed non-random interactions with the heterogeneous membrane; however, both the magnitude and concentration dependence of the apparent preference differed markedly between compounds (Fig. 1C). Representative top-view snapshots at a drug-to-lipid ratio of 0.10 are shown in Fig. S2, while the corresponding number-density profiles across the bilayer are provided in Fig. S3. CHX showed a persistent preference for the domain-associated lipid environment across the entire investigated concentration range. ΔΔ*G_app_* changed from -3.55 kJ mol*^−^*^1^ at 0.05 drug/lipid to -3.06 kJ mol*^−^*^1^ at 0.1 and -1.28 kJ mol*^−^*^1^ at 0.3. Although the magnitude of the preference decreased with increasing CHX loading, preferential association was retained even at the highest investigated concentration. The lipid-resolved analysis indicated that this preference was particularly associated with PG-rich local environments, whereas PE was consistently depleted around CHX (Fig. S4). This observation is compatible with the known ability of CHX to interact with phospho-lipid headgroups; however, it should not be reduced to a simple electrostatic mechanism. Neutron diffraction localized CHX near the glycerol backbone of phosphatidylcholine bilayers, consistent with an interfacial, wedge-like configuration.^32^ Molecular simulations and NMR experiments using *S. aureus*- and *E. coli* -like membrane models similarly placed CHX at the lipid headgroup–tail interface and showed that its partitioning depends on membrane composition.^33^ Yet substantial CHX uptake into homogeneous DMPC bilayers was accompanied by only modest changes in bulk membrane structure.^34^ Together with our previous findings, CHX does not exhibit simple selectivity for globally anionic membranes, ^6^ the present results are consistent with local membrane composition and physicochemical environment contributing to CHX association beyond membrane-averaged charge alone. The present results extend this view by suggesting that, in a laterally heterogeneous membrane, CHX preferentially samples a specific local compositional environment rather than simply discriminating membranes according to their net surface charge.

OCT showed a different concentration dependence. At the lowest loading, its apparent preference free energy was *−*0.94 kJ mol*^−^*^1^, indicating preferential association with the domain-associated environment. A similar, slightly stronger preference was observed at 0.1 drug/lipid, where ΔΔ*G*_app_ = *−*1.18 kJ mol*^−^*^1^, whereas the preference was largely lost at the highest loading, with ΔΔ*G*_app_ = *−*0.29 kJ mol*^−^*^1^. Thus, OCT displayed a modest preference for the domain-associated environment at low and intermediate loading, which approached near-neutrality at the highest drug/lipid ratio. This concentration dependence is consistent with the collective behavior previously observed for OCT. In our earlier simulations, OCT formed aggregates before interaction with the membrane, whereas CHX remained considerably more dispersed and interacted predominantly as individual molecules.^6^ Consequently, increasing OCT loading may progressively shift the balance from lipid-environment-dependent interactions toward OCT–OCT interactions and collective surface association. This interpretation also follows previous experimental observations by Malanovic et al., who showed that OCT inserts into the acyl-chain region and induces pronounced lipid disorder, with hydrophobic interactions becoming increasingly important in membrane disruption.^7^ Time-resolved SAXS measurements on PE/PG/CL vesicles have since revealed an OCT-induced sequence of membrane lysis and lipid reassembly into multilamellar structures, accompanied by changes in bilayer thickness and the gel-to-fluid transition.^35^ These observations provide independent experimental evidence that increasing OCT exposure can drive the membrane from local bilayer perturbation toward large-scale structural reorganization. Therefore, the loss of apparent compositional preference at high OCT loading does not necessarily indicate weaker membrane interaction. Instead, it may reflect a transition from an initially selective interaction toward a more collective and less composition-dependent mode of membrane association.

PmB exhibited the strongest apparent preference among the three antimicrobial agents. In contrast to OCT and CHX, a substantial fraction of PmB molecules remained in the aqueous phase rather than becoming membrane-associated. This behaviour is already evident at a drug-to-lipid ratio of 0.10, where the PmB density extends prominently into the aqueous region. By contrast, OCT and CHX show broader antimicrobial density distributions at the highest loading, consistent with the formation of larger drug aggregates and an increasing contribution from drug–drug interactions (Fig. S3). For the membrane-associated population, ΔΔ*G*_app_ was *−*5.08, *−*2.32, and *−*0.26 kJ mol*^−^*^1^ at drug/lipid ratios of 0.05, 0.1, and 0.3, respectively. Thus, membrane-associated PmB showed a pronounced preference for the domain-associated lipid environment at low loading, but this preference weakened markedly with increasing PmB concentration and approached near-neutrality at the highest loading. The lipid-resolved analysis further indicated that this effect was strongly associated with HOP-rich and PE-depleted local environments (Fig. S4). Since HOP promotes ordered domain formation in the present membrane model, its enrichment around PmB suggests preferential association with a hopanoid-enriched lateral environment rather than a specific PmB–HOP interaction. Diplopterol has previously been shown to modulate membrane affinity, poration activity, and the time-dependent mechanical response to the antimicrobial peptide Polybia-MP1.^36^ Although Polybia-MP1 is chemically distinct from PmB, these findings provide direct evidence that hopanoids can influence antimicrobial– membrane interactions beyond membrane charge alone. The preferential HOP enrichment observed here therefore supports sensitivity of PmB to the organized lipid environment rather than evidence for a specific PmB–hopanoid binding interaction. The strong dependence of PmB interactions on membrane composition is consistent with previous simulation studies. Berglund et al. showed that PmB interacts differently with chemically distinct bacterial membrane environments and that membrane composition substantially determines peptide binding and insertion behavior.^37^ Similarly, coarse-grained simulations have shown that PmB can remain shallowly associated with phospholipid membranes and strongly influence their physical properties without necessarily producing immediate membrane permeabilization. ^14^ Our results further suggest that lateral organization within a single heterogeneous membrane may provide an additional level of selectivity.

Taken together, the apparent preference free energies demonstrate that the three antimicrobial agents do not interact with the heterogeneous membrane in an equivalent manner. CHX maintained a clear preference for the domain-associated environment across the investigated loading range, OCT showed a modest preference at low and intermediate loading that was largely lost at the highest loading, whereas the membrane-associated PmB population displayed the strongest initial preference but also the largest loading-dependent decrease. These results suggest that preferential interaction with lateral membrane organization may represent a common feature of antimicrobial association, while the strength and concentration dependence of this interaction remain compound-specific.

### Preferential association is accompanied by compound-specific changes in lipid mobility

We next investigated whether the observed differences in local membrane association were accompanied by changes in membrane dynamics. Among the determined structural and dynamic parameters, lateral diffusion showed the clearest concentration-dependent response and was therefore used as the principal descriptor of lipid mobility. Diffusion is not interchangeable with membrane fluidity or phase state, but it is a sensitive reporter of heterogeneous antimicrobial-membrane interactions. Fluorescence correlation spectroscopy of magainin 2 and mastoparan X, for example, resolved peptide-induced subpopulations of lipids with altered diffusion even at low peptide-to-lipid ratios.^38^ We therefore interpret the diffusion coefficients below as measures of molecular mobility and dynamic heterogeneity rather than as a complete description of membrane material state (Fig. 2).

**Figure 2:**
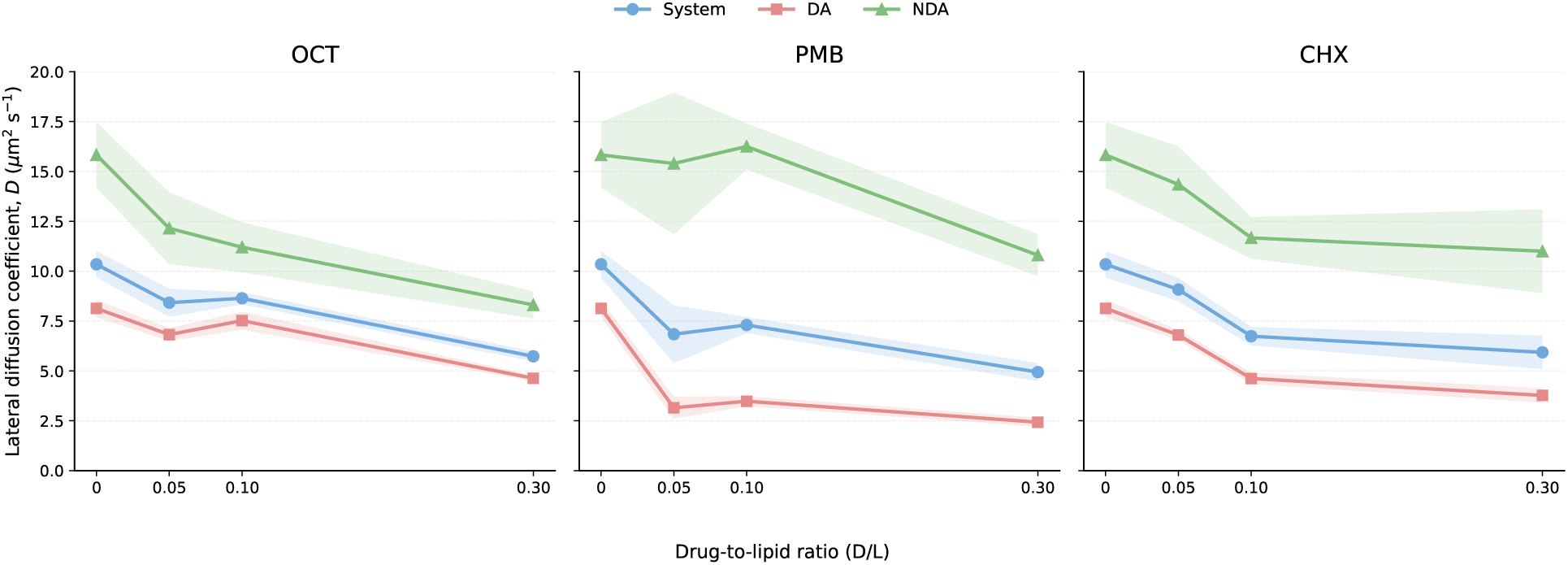
Preferential association is accompanied by compound-specific changes in membrane dynamics. Lateral lipid diffusion coefficients for OCT (left), PmB (centre), and CHX (right) at drug-to-lipid ratios of 0.05, 0.10, and 0.30. Values are shown for the whole membrane and separately for the domain-associated and non-domain-associated lipid pools. Symbols and connecting lines show mean values, while shaded regions indicate temporal variability over the analysed trajectory interval.

CHX induced a progressive reduction in lipid diffusion with increasing loading. The system-averaged diffusion coefficient decreased from 9.1 to 5.9 *µ*m^2^ s*^−^*^1^ between the lowest and highest investigated concentrations. A stronger reduction was observed for the domain-associated lipid pool, for which diffusion decreased from 6.8 to 3.8 *µ*m^2^ s*^−^*^1^. Although the non-domain-associated lipid population remained more mobile, its diffusion was also progressively reduced. The decrease in mobility therefore followed the persistent apparent preference of CHX for the domain-associated environment. This suggests that CHX association is accompanied predominantly by restriction of local lipid dynamics rather than by extensive structural reorganization of the entire bilayer. Such a result follows previous descriptions of CHX activity, in which lower concentrations of CHX were associated with reduced membrane fluidity and impairment of membrane-dependent cellular processes.^6^ The present model additionally suggests that this effect can be enhanced within the lateral environment preferentially sampled by CHX.

The concentration dependence of OCT was different. Lipid mobility remained relatively similar between the two lower OCT loadings, whereas a pronounced decrease was observed at 0.3 drug/lipid. System-averaged diffusion decreased from 8.4 to 5.7 *µ*m^2^ s*^−^*^1^, and both domain-associated and non-domain-associated lipid populations showed reduced mobility at the highest loading. Interestingly, the strongest reduction in diffusion appeared in the same concentration range in which the apparent compositional preference of OCT approached zero. This suggests that increasing OCT concentration may alter the character of its membrane interaction. At low loading, OCT preferentially samples the domain-associated environment, whereas at higher loading its interaction becomes less composition-selective but affects a broader fraction of the membrane. Such a transition is consistent with the tendency of OCT to form supramolecular aggregates and with previous reports of widespread OCT-induced disordering of membrane lipids.^7,20^ A functional counterpart has recently been observed in intact purple bacteria, where low-micromolar OCT strongly impaired membrane energization and cytochrome turnover while leaving primary reaction-centre photochemistry comparatively preserved.^39^ As these processes rely on the spatial organization and mobility of membrane-associated redox components, the results are consistent with membrane-level reorganization compromising collective membrane function before complete membrane disintegration.

PmB also showed a pronounced decrease in lipid mobility at the highest investigated loading. The system-averaged diffusion coefficient decreased from 6.9–7.3 *µ*m^2^ s*^−^*^1^ at the two lower concentrations to 4.9 *µ*m^2^ s*^−^*^1^ at 0.3 drug/lipid. The domain-associated lipid population remained substantially less mobile than the non-domain-associated population throughout the simulations and showed a further decrease at the highest PmB loading. Previous simulations of PmB interacting with bacterial phospholipid membranes have similarly shown reduced lateral lipid diffusion following peptide association. Berglund et al. reported restricted lipid movement following PmB binding, while later coarse-grained simulations showed that PmB can suppress phospholipid mobility without inducing extensive membrane disruption.^37^ In a more recent, protein-rich model of the *E. coli* outer membrane, PmB formed bridges between LPS and outer-membrane proteins and reduced the lateral diffusion of both components by approximately two-to threefold.^40^ The present results follow these observations and suggest that even a membrane-associated subpopulation of PmB can influence the dynamic state of the heterogeneous bilayer. The agreement across phospholipid, LPS-containing, and protein-containing models also indicates that restricted lateral mobility is a robust consequence of PmB association, although the molecular species immobilized in each membrane are different. In contrast, global APL and membrane thickness showed substantially weaker and less systematic concentration-dependent changes for all three compounds (Fig. S5). These parameters were therefore not considered primary descriptors of the antimicrobial response in the present analysis. Their relative stability further suggests that the changes in membrane dynamics cannot be explained by a simple uniform expansion, compression, or thinning of the bilayer.

### Changes in lipid dynamics are accompanied by distinct mechanical responses

The different modes of antimicrobial association were also reflected in the mechanical properties of the membrane. For CHX, bending rigidity was reduced relative to the untreated membrane (*κ* = 40.8 *k*_B_*T*). but remained comparatively stable across the investigated loading range, varying between approximately 28 and 31 *k*_B_*T* . Although CHX substantially decreased lipid diffusion, this dynamic restriction was not accompanied by a comparably strong concentration-dependent change in global membrane elasticity. This behavior suggests that CHX predominantly modifies local membrane dynamics and organization rather than inducing extensive mechanical destabilization of the whole bilayer. In our previous work on homogeneous membranes, CHX similarly produced discrete changes in membrane properties without causing membrane disruption as its primary activity.^6^ Coarse-grained simulations of several cationic antiseptics also found compound-specific dynamic and structural perturbations that sensitized bacterial-like bilayers to electric-field-induced poration even when the membranes did not spontaneously disintegrate.^27^ Such sensitization illustrates how a local change in organization can alter membrane susceptibility without a large monotonic shift in a single global elastic parameter. Our earlier study also raised the possibility that CHX-induced physical changes could interfere with the ability of bacterial membranes to maintain lateral microdomain organization, although this hypothesis could not be addressed directly because the investigated systems were homogeneous.^6^ The present heterogeneous model therefore provides a more direct basis for examining this possibility.

OCT produced a substantially different mechanical response. Bending rigidity progressively decreased from 26 to 23 *k*_B_*T* . At the highest loading, this response was accompanied by a pronounced increase in interdigitation (Fig. S6). The combined behavior of bending rigidity and interdigitation may provide a more informative description of membrane remodeling than any individual structural parameter alone. In particular, the simultaneous reduction in bending indicates an increased susceptibility of the membrane to deformation, while increased interdigitation suggests substantial rearrangement of interleaflet packing. Together with the reduction in lipid mobility at high loading, the data support a concentration-dependent transition toward broader mechanical remodeling of the membrane. This follows previous observations of OCT-induced disruption of lipid packing and membrane integrity. Malanovic et al. reported that OCT penetrates into the hydrophobic membrane region and induces strong lipid disorder and changes in membrane fluidity. ^7^ Our previous work also showed that the membrane response to OCT could not be explained solely by simple electrostatic selectivity and suggested an important contribution from membrane mechanical properties.^6^ The present results further indicate that these mechanical effects become increasingly pronounced as the initial compositional preference of OCT decreases.

PmB showed an opposite mechanical tendency. Bending rigidity increased from 32 *k*_B_*T* at the lower loading levels to 35 *k*_B_*T* at 0.3 drug/lipid. Interdigitation did not show a comparable systematic increase. This response indicates that the restriction of lipid mobility induced by PmB is accompanied by an increase in membrane mechanical resistance rather than by OCT-like softening. Our results follow previous coarse-grained simulations showing that PmB can stiffen phospholipid bilayers, increase membrane bending rigidity, and restrict lipid diffusion.^14^ In those studies, the effect was attributed to PmB filling lipid packing defects and increasing lipid order. High-resolution AFM of native *E. coli* outer-membrane patches has likewise shown that polymyxins organize LPS into crystalline assemblies that increase membrane stiffness while decreasing thickness and increasing area. ^41^ While the detailed structural mechanism in the present hopanoid-containing membrane differs from both a simple phospholipid bilayer and an LPS outer membrane, these results collectively show that PmB can mechanically constrain one membrane environment while destabilizing another. The direction of the elastic response is therefore an emergent property of membrane composition and lateral organization rather than a fixed signature of PmB binding.

### Molecular-scale membrane remodeling is reflected in vesicle-scale softening

Having established that the investigated antimicrobials differ in their association with lateral lipid environments and in their effects on lipid dynamics, we next examined whether these molecular-scale changes are reflected in the mechanical response of the membrane. Membrane bending rigidity provides a collective measure of the energetic cost of membrane deformation and is therefore sensitive to changes in lipid packing, organization, and intermolecular interactions.^25,42^ Antimicrobial-induced softening is not unique to the present compounds: fluctuation analysis showed a pronounced concentration-dependent decrease in the bending modulus of POPC GUVs exposed to magainin 2. ^43^ At the same time, experiments with Polybia-MP1 demonstrated that a diplopterol-containing membrane can soften initially and recover stiffness at longer times.^36^ Together, these studies support membrane elasticity as a sensitive readout of antimicrobial action while highlighting the importance of interpreting mechanical changes in the context of membrane composition, exposure history, and other dynamic observables. We used fluctuation spectroscopy of GUVs to determine whether the compound-dependent membrane perturbations identified in the simulations are accompanied by changes in membrane mechanics at the vesicle scale (Fig. 3).

**Figure 3:**
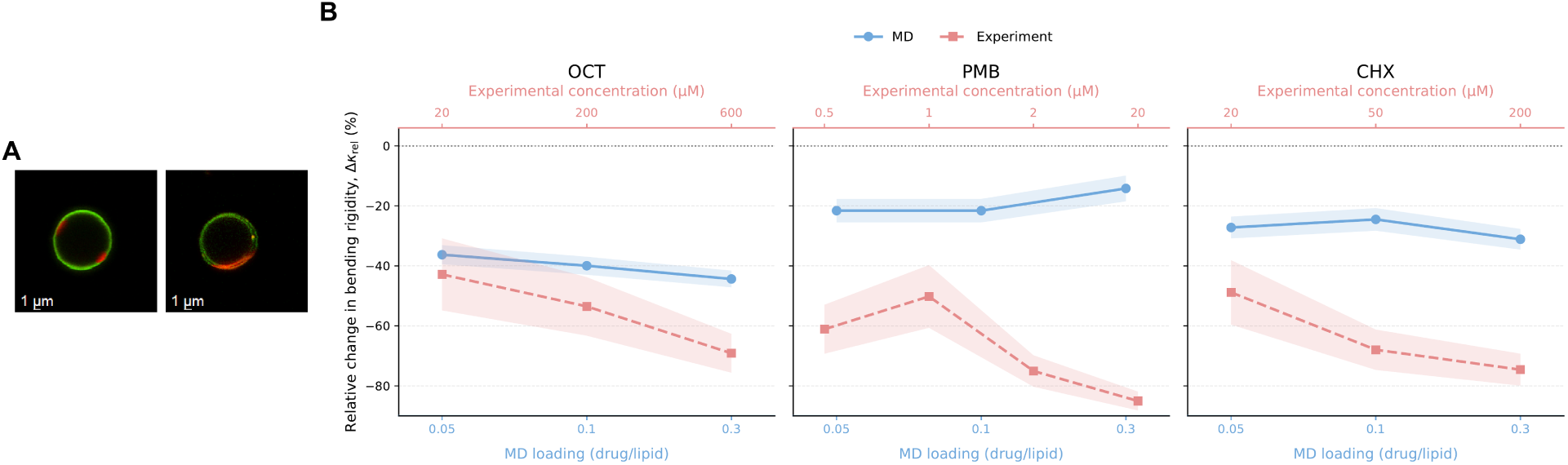
Molecular-scale membrane remodeling is reflected in vesicle-scale membrane softening. (A) Representative dual-channel confocal images of individual GUVs used for fluctuation analysis showing Atto488-DOPE in green and DiIC_12_ in red. The spatially heterogeneous distribution of the probes visualizes lateral membrane heterogeneity (B) Relative changes in bending rigidity, Δ*κ*_rel_, obtained from MD simulations (blue circles; lower x-axes) and GUV fluctuation spectroscopy (red squares; upper x-axes) for OCT, PmB, and CHX. Simulation conditions are expressed as drug-to-lipid ratios, whereas experimental conditions are expressed as bulk antimicrobial concentrations.

All three compounds produced a pronounced decrease in bending rigidity relative to the untreated membrane. OCT reduced *κ* by 42.8, 53.5, and 69.1% at 20, 200, and 600 *µ*M, respectively, showing a progressive softening with increasing concentration. CHX produced an even stronger response, with reductions of 48.8, 68.0, and 74.6% over the investigated concentration range. PmB affected membrane mechanics already at considerably lower concentrations; the bending rigidity was reduced by 61.1 and 50.2% at 0.5 and 1 *µ*M, respectively, followed by decreases of 75.0 and 85.0% at 2 and 20 *µ*M. Thus, despite differences in the concentration dependence, the experimental measurements consistently showed that antimicrobial exposure shifts the heterogeneous membrane toward a more deformable state. The same direction of the mechanical response was observed independently in the MD simulations. Relative to the untreated membrane, OCT decreased the bending rigidity by 36.3–44.4% over the investigated drug-to-lipid ratios, with the magnitude of softening increasing with loading. CHX produced a reduction of approximately 25–31%, while PmB decreased *κ* by approximately 14–22%. The magnitude of the experimental response was therefore larger, but the two approaches followed the same overall tendency: each antimicrobial reduced the resistance of the membrane to bending. More importantly, the additional molecular descriptors obtained from MD indicate that this common mechanical outcome is associated with different patterns of membrane reorganization. The corresponding absolute bending rigidities and the diameter distributions of the GUV populations analysed experimentally are provided in Fig. S7 and Fig. S8, respectively.

For OCT, the mechanical response was particularly consistent across the two scales. The progressive decrease in experimental *κ* followed the loading-dependent softening observed in MD. At the molecular level, this response developed together with a gradual loss of the initial preference of OCT for domain-associated lipid environments, reduced lipid mobility at the highest loading and increased interdigitation. Taken together, these changes support a transition from preferential association at low loading toward increasingly collective membrane remodeling as the amount of OCT in the membrane increases. This interpretation follows our previous observations that OCT activity is strongly related to membrane mechanical properties and cannot be explained solely by electrostatic interactions.^6^ In the present heterogeneous membrane system, the simulations further show that the mechanical response is accompanied by a change in how OCT samples different lateral lipid environments. Previous work on OCT and CHX similarly treated bending rigidity together with other structural and dynamic observables rather than as an isolated parameter.^6^

CHX produced a different molecular response despite leading to the same overall direction of membrane softening. Its preference for domain-associated lipid environments remained detectable throughout the investigated loading range, while lipid diffusion progressively decreased, particularly within the domain-associated lipid pool. In MD, the initial reduction in bending rigidity was followed by relatively modest changes with further loading, whereas the GUV measurements showed progressively stronger softening with increasing CHX concentration. This difference may indicate that the local perturbations identified in the simulations, including restricted lipid mobility and persistent preferential association, become more pronounced when expressed over the larger spatial and temporal scales of the vesicle membrane. Previous studies have shown that CHX can alter lipid organization and membrane mechanical properties without requiring immediate membrane disintegration. ^6^ Our earlier work also proposed that changes in membrane stiffness could interfere with lateral membrane organization, although the homogeneous membrane models used at that time did not permit this possibility to be examined directly. The present simulations provide a molecular view of this response by linking CHX-induced softening with persistent interactions with specific lateral lipid environments and a substantial reduction in lipid mobility.

PmB showed the strongest experimental response at comparatively low concentrations. Although the concentration dependence was not fully monotonic at the two lowest concentrations, bending rigidity decreased markedly above 1 *µ*M and reached an approximately 85% reduction at 20 *µ*M. The simulations also showed lower bending rigidity than in the untreated membrane, but the response was weaker and partially recovered at the highest loading. At the molecular level, PmB differed from both OCT and CHX because the membrane-associated population displayed a strong preference for domain-associated environments and was accompanied by reduced lipid mobility at higher loading. Previous simulations of PmB have demonstrated that its influence on membrane mechanics depends strongly on membrane composition: PmB was reported to restrict lipid diffusion and increase bending rigidity in a phospholipid inner-membrane model, whereas studies of asymmetric Gram-negative membranes showed disruption of lipid homeostasis, leaflet stress imbalance, lipid scrambling, phase separation, and formation of defects at lateral domain boundaries.^14,15^ Together with the LPS crystallization observed experimentally in native outer-membrane patches,^41^ these observations support the view that the mechanical response to PmB depends strongly on the organization of the local lipid environment rather than being an intrinsic consequence of PmB binding alone. The pronounced PmB response at low experimental concentrations is also consistent with its high antibacterial potency. PmB activity against susceptible Gram-negative isolates is commonly reported in the low mg L*^−^*^1^ range,^24^ placing the low-micromolar concentrations used here within a biologically relevant exposure regime. This point is important when comparing the concentration-dependent responses obtained from the different approaches. The simulations are defined by membrane drug-to-lipid loading, whereas the experiments are defined by the antimicrobial concentration in the surrounding solution; consequently, the corresponding exposure levels should be interpreted as relative concentration series rather than as directly equivalent conditions. This distinction is particularly relevant for compounds such as PmB, for which only a fraction of the molecules becomes membrane-associated.

Overall, the bending measurements show that the local antimicrobial-induced perturbations identified at the molecular level are accompanied by a substantial change in the collective mechanical behaviour of the membrane. Although OCT, CHX, and PmB all shifted the membrane toward lower bending rigidity, the simulations indicate that this common vesicle-scale response arises together with markedly different patterns of lateral association, lipid mobility, and membrane remodeling. Fluctuation spectroscopy therefore provides an independent readout of the resulting membrane mechanical state, while MD allows this response to be interpreted in relation to the heterogeneous lipid environments sampled by each compound. Bending rigidity thus captures the collective mechanical consequence of the local membrane perturbations resolved in the simulations, integrating their effects across the heterogeneous GUV membrane. We therefore next examined whether this altered mechanical state translates into changes in whole-vesicle stability under confinement using the microfluidic trapping assay.

### Antimicrobial-induced membrane remodeling alters whole-vesicle mechanical stability

The molecular and vesicle-scale measurements showed that antimicrobial association with the heterogeneous membrane is accompanied by changes in lipid dynamics and a decrease in bending rigidity. We therefore examined whether these changes are also reflected in the ability of the membrane to maintain whole-vesicle stability under confinement. GUVs were retained in microfluidic traps and exposed to the investigated compounds under continuous flow, while the fraction of vesicles remaining in the traps was followed over time. Microfluidic confinement provides a useful means of probing membrane behaviour under controlled mechanical conditions, as retention and release of GUVs depend on the balance between hydrodynamic forces, vesicle deformation, and the elastic response of the membrane. ^26,44^ Related microfluidic studies have shown that GUV trapping can remain stable during solution exchange and flow, while changes in membrane deformability alter the response of vesicles to confinement.^45,46^ High-throughput single-GUV measurements under continuous antimicrobial-peptide dosing have shown that both peptide concentration and mode of action are reflected in the timing and distribution of disruption events across vesicle populations.^47^ These findings support population-level retention kinetics as a sensitive functional readout, while emphasizing that the resulting curves integrate heterogeneous responses of individual vesicles (Fig. 4).

**Figure 4:**
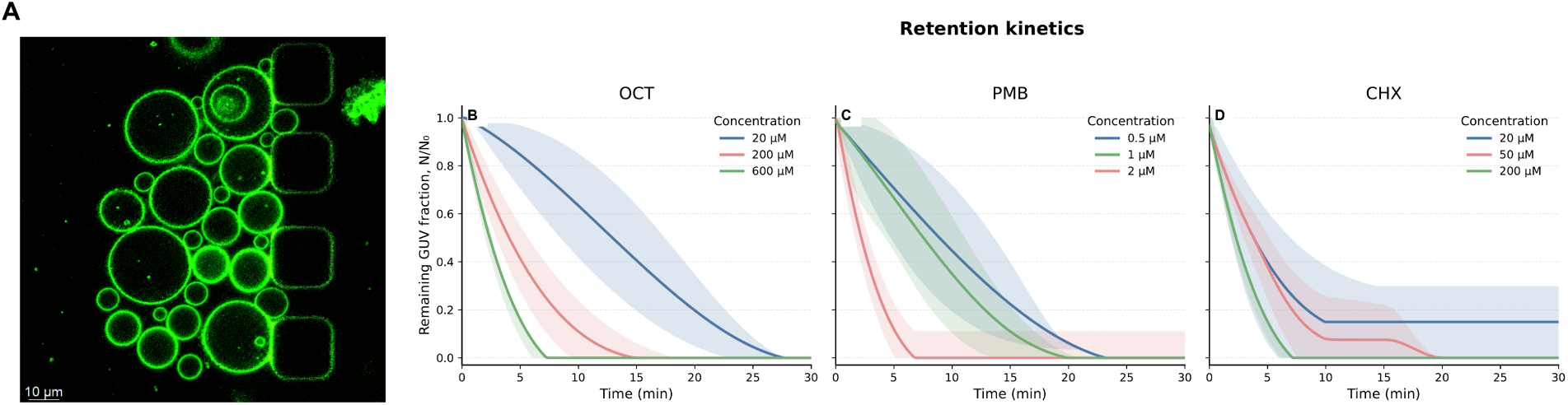
Antimicrobial-induced membrane remodeling alters whole-vesicle mechanical stability. (A) Representative confocal image of GUVs retained in the microfluidic trapping structures. Fluorescence was collected in a single acquisition channel and is displayed in green. Remaining GUV fractions, *N/N*_0_, during continuous exposure to (B) OCT, (C) PmB, and (D) CHX at the indicated bulk concentrations. Solid lines show the mean retention kinetics, while shaded regions indicate variability between independent experiments.

All three antimicrobials produced a concentration-dependent acceleration of GUV loss from the traps (Fig. 4). This behaviour was particularly clear for OCT, for which the characteristic time required for the retained population to decrease to approximately one half shifted from about 13 min at 20 *µ*M to approximately 4 min at 200 *µ*M and 2.4 min at 600 *µ*M. A similar response was observed for PmB despite the substantially lower concentrations required: the approximate half-retention times were 8.9, 7.5, and 2.2 min for 0.5, 1, and 2 *µ*M, respectively. CHX also showed accelerated vesicle loss, with the corresponding values decreasing from approximately 3.8 min at 20 *µ*M to 2.3 min at 50 *µ*M. These values are used here only as descriptive measures of the kinetic shift; the full retention curves provide the primary comparison between conditions.

For OCT, the retention kinetics closely follow the sequence of changes observed at the preceding levels of membrane organization. Increasing OCT loading in MD was accompanied by a gradual loss of its initial preference for domain-associated lipid environments and increasingly pronounced changes in lipid mobility, interdigitation and bending rigidity. At the GUV scale, bending rigidity similarly decreased with increasing OCT concentration, while the microfluidic measurements now show that this response is accompanied by progressively faster loss of whole-vesicle stability. These results follow our previous observations in which OCT activity was associated more strongly with the mechanical state of the membrane than with simple electrostatic selectivity. ^6^ In that earlier study, changes in bending rigidity and membrane disruption were also concentration dependent, although the homogeneous membrane models did not allow the role of lateral heterogeneity to be addressed directly.^6^ The present results extend this picture by linking OCT-induced changes in membrane mechanics with its evolving association with different lateral lipid environments.

CHX reached the same macroscopic outcome through a different molecular response. In contrast to OCT, CHX retained a preference for domain-associated environments over the investigated loading range and progressively restricted lipid mobility, while the MD bending response changed comparatively little after the initial softening. At the vesicle level, how-ever, increasing CHX concentration produced both stronger membrane softening and faster GUV loss. This indicates that restricted molecular mobility and persistent local association can coexist with increasing susceptibility of the membrane as a whole to deformation under confinement. Previous studies have shown that CHX can substantially modify lipid organization and membrane mechanical properties without requiring immediate catastrophic permeabilization.^6^ Electron microscopy of intact *B. subtilis* and *E. coli* also revealed time- and concentration-dependent CHX-induced indentations whose spatial distributions differed between the two organisms, leading the authors to propose a contribution from the non-uniform distribution of membrane lipids.^48^ Although cell-wall architecture precludes a direct mapping to our GUVs, these observations are consistent with spatially heterogeneous membrane damage rather than a purely uniform detergent-like response. This distinction is relevant here: the microfluidic response does not require that CHX acts through the same local process as OCT, but shows that their different molecular-scale interactions ultimately affect the ability of the heterogeneous membrane to sustain an imposed mechanical challenge.

PmB provides perhaps the clearest indication that whole-vesicle stability integrates several aspects of the membrane response rather than bending rigidity alone. Experimental bending rigidity was not fully monotonic between 0.5 and 1 *µ*M PmB, whereas GUV retention showed a clearer acceleration with increasing concentration, becoming particularly rapid at 2 *µ*M. In the simulations, the membrane-associated PmB population displayed the strongest preference for domain-associated lipid environments and restricted lipid diffusion at higher loading. Previous computational work similarly showed that PmB can strongly perturb lipid diffusion and membrane mechanics, with the direction and magnitude of the response depending on membrane composition and organization.^14^ Thus, the pronounced response observed here at low PmB concentrations may reflect the combined consequences of selective membrane association, changes in lateral organization, and altered membrane mechanics rather than a simple dependence on a single elastic coefficient.

Such coupling between lateral membrane organization and antimicrobial activity has been proposed more broadly for cationic membrane-active compounds. Antimicrobial molecules can redistribute anionic lipids and promote lateral lipid segregation, thereby modifying membrane organization beyond their immediate binding sites.^8,49^ This is particularly relevant for heterogeneous bacterial membranes, where lipid domains themselves contribute to spatial organization and membrane function.^2,50^ Our results suggest that such local interactions may propagate into changes in collective membrane properties and ultimately alter the ability of the entire vesicle to withstand deformation.

Comparison across the three compounds shows that the strength of local compositional preference does not directly predict the subsequent mechanical response. PmB retained the strongest preference for domain-associated environments, yet its MD bending response partially recovered at the highest loading. OCT showed the opposite tendency: its initial compositional preference weakened with increasing loading, while changes in interdigitation and vesicle stability became more pronounced. CHX, in turn, maintained preferential association while the experimental mechanical response continued to increase. These differences suggest that local recruitment and membrane-wide destabilization are related, but not simply coupled. Preferential environments may favor the initial accumulation of a compound, whereas the later response is likely shaped by additional, compound-specific processes, including aggregation, saturation of preferred regions, and the spread of local perturbations through the surrounding membrane. In this sense, compositional selectivity and the resulting material response should be considered together rather than treating either as a direct proxy for antimicrobial potency.

The responses resolved here represent the passive material behavior of the membrane rather than the full cellular response to antimicrobial exposure. Lipid-only GUVs isolate this contribution, while omitting leaflet asymmetry, LPS, membrane proteins, the cell wall, and active homeostatic processes. Even so, direct comparison of antimicrobial-peptide action on GUVs and *E. coli* spheroplasts has shown that simplified and bacterial membranes can reach quantitatively similar steady-state permeabilities despite their structural differences. ^51^ At the same time, recent live-cell work showed that PmB killing at clinically relevant concentrations involves energy-dependent remodeling of the outer membrane, including LPS loss and surface protrusion, before subsequent access to and permeabilization of the inner membrane.^52^ The retention kinetics measured here should therefore not be read as direct bactericidal kinetics, but as a passive mechanical component of antimicrobial action that, in intact cells, operates together with envelope structure and active membrane remodeling.

Overall, the microfluidic experiments extend the analysis to the behaviour of the entire vesicle under confinement. Increasing antimicrobial exposure consistently accelerated GUV loss from the trapping structures, demonstrating that the effects of OCT, CHX, and PmB extend beyond local lipid interactions and collective membrane elasticity to the vesicle-scale response. Importantly, the microfluidic readout integrates multiple properties of the treated membrane rather than reporting a single molecular or mechanical parameter. Loss of a trapped vesicle may result either from membrane rupture or from sufficient deformation to allow the vesicle to escape the trapping structure; in the present context, both outcomes indicate an altered response of the antimicrobial-treated vesicle to confinement. Considered together with the MD and fluctuation-spectroscopy results, these measurements support a multiscale picture in which compound-specific interactions with laterally organized lipid environments are accompanied by changes in lipid dynamics, membrane mechanics, and whole-vesicle behaviour. Rather than defining a single linear sequence, these complementary readouts show how distinct molecular responses can converge at the vesicle scale.

The compound-specific responses observed here also motivate future studies of combined antimicrobial exposure. Such experiments could test whether changes induced by one compound alter the membrane association or mechanical effects of another. The present comparison of individual compounds provides a basis for these studies.

Lateral membrane heterogeneity creates distinct physicochemical environments that shape antimicrobial association and the subsequent membrane response. By combining coarsegrained molecular dynamics, GUV fluctuation spectroscopy, and microfluidic trapping, we followed this response from local antimicrobial-lipid interactions to collective membrane mechanics and whole-vesicle stability. OCT, CHX, and PmB all showed preferential association with domain-associated membrane environments, although the strength and loading dependence of this preference were compound-specific. CHX retained a clear preference across the investigated range, OCT showed only modest preference at low and intermediate loading that was largely lost at the highest loading, whereas PmB displayed the strongest initial preference followed by a pronounced reduction with increasing loading.

These distinct association patterns were accompanied by compound-specific changes in lipid dynamics, yet all three antimicrobials ultimately shifted the membrane toward lower bending rigidity. This tendency was observed in both simulations and GUV fluctuation spectroscopy, linking molecular-scale perturbations to the collective elastic response of the membrane. At the vesicle scale, increasing antimicrobial exposure also accelerated GUV loss from microfluidic traps, indicating a progressive reduction in whole-vesicle mechanical stability.

Taken together, our results show that a common membrane-level outcome does not require a common molecular pathway. Different patterns of local association and lipid remodeling can converge on membrane softening and reduced mechanical stability. More broadly, these findings identify lateral membrane heterogeneity as an active determinant of antimicrobial response and show that membrane-active compounds cannot be understood solely from bulk lipid composition or membrane-averaged charge. Instead, the spatial organization of membrane components provides an additional level of control over antimicrobial association and its propagation toward collective membrane behaviour.

## Author Contributions

MR conceived the study and designed the research. MR and DD developed the experimental membrane model and performed the GUV imaging and membrane fluctuation measurements. NY designed and developed the microfluidic trapping platform, provided the microfluidic devices and experimental protocol, and contributed to the implementation of the GUV trapping experiments. MR performed the molecular dynamics simulations and computational analyses. MR and DD analyzed the experimental data. MR analyzed simulation data and prepared the figures. MR supervised the project and contributed to the interpretation of the results. MR wrote the original draft of the manuscript. All authors contributed to the discussion of the results, reviewed and edited the manuscript, and approved the final version.

## Acknowledgments

M.R. thanks to the National Science Centre, Poland (grant no. 2022/45/N/NZ9/02130) for financial support. The authors gratefully acknowledge Maxime Kucharski from Wroc-law University of Science and Technology for technical assistance with GUV electroformation.

## Data Availability Statement

The datasets used in this study can be found in the Zenodo repository.

