## Supplementary Materials for "Lateral Membrane Heterogeneity Shapes Antimicrobial Association and Convergent Mechanical Responses"

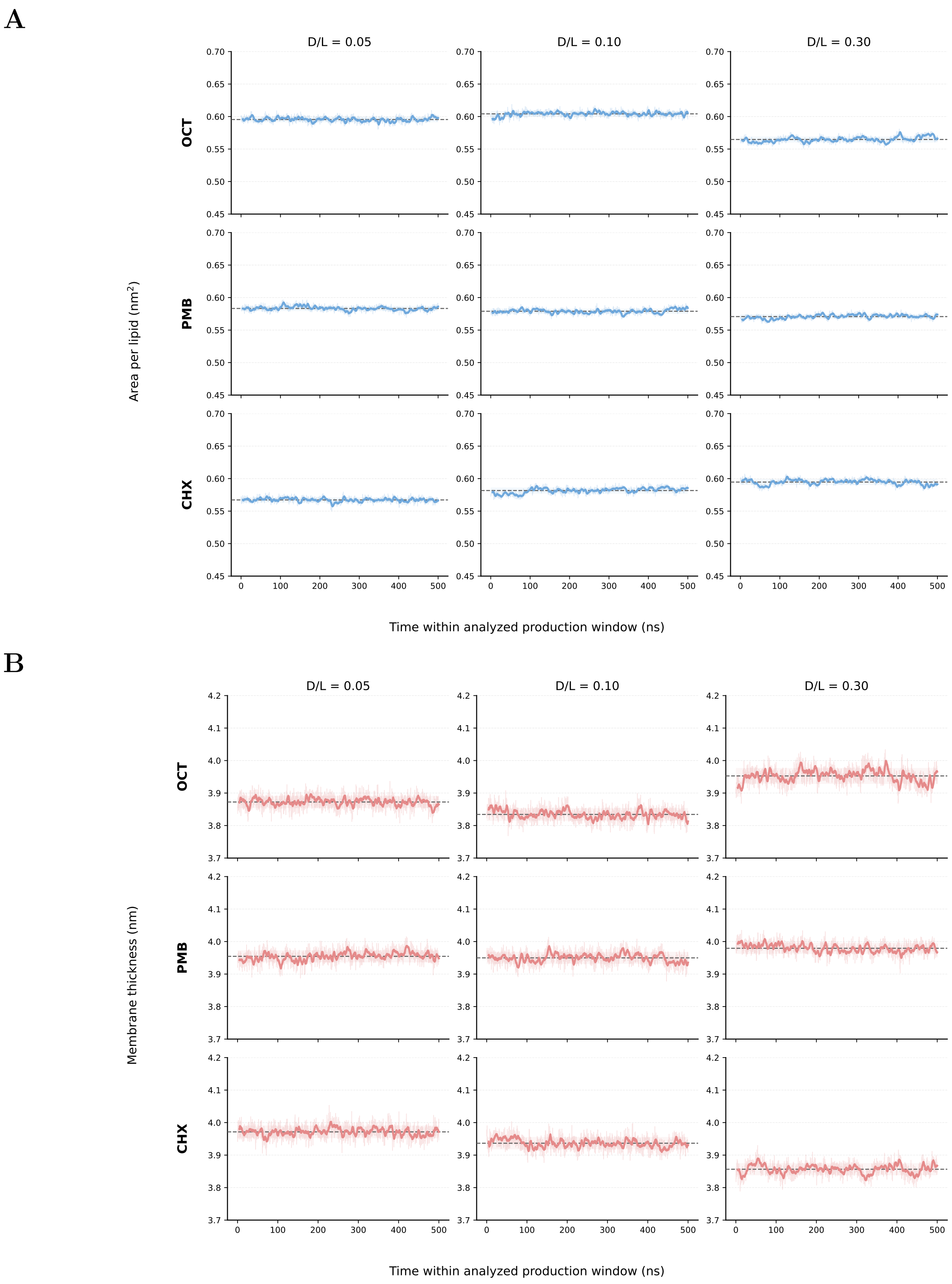


Figure 1: Convergence of the membrane structural parameters over the analysed production interval. (A) Area per lipid and (B) membrane thickness as a function of simulation time for OCT, PmB, and CHX at drug-to-lipid ratios (D/L) of 0.05, 0.10, and 0.30. Dashed horizontal lines indicate the mean values over the final 500 ns used for quantitative analysis.

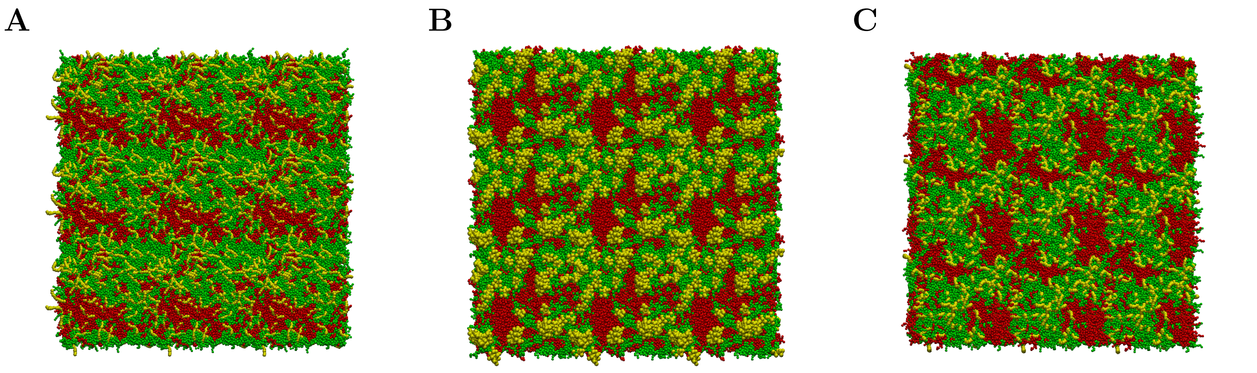


Figure 2: Representative top-view snapshots of the simulated heterogeneous membrane at a drug-to-lipid ratio of 0.10 for (A) OCT, (B) PmB, and (C) CHX. Domain-associated and non-domain-associated lipid pools are shown in red and green, respectively, and antimicrobial molecules are shown in yellow. Water and ions are omitted for clarity.

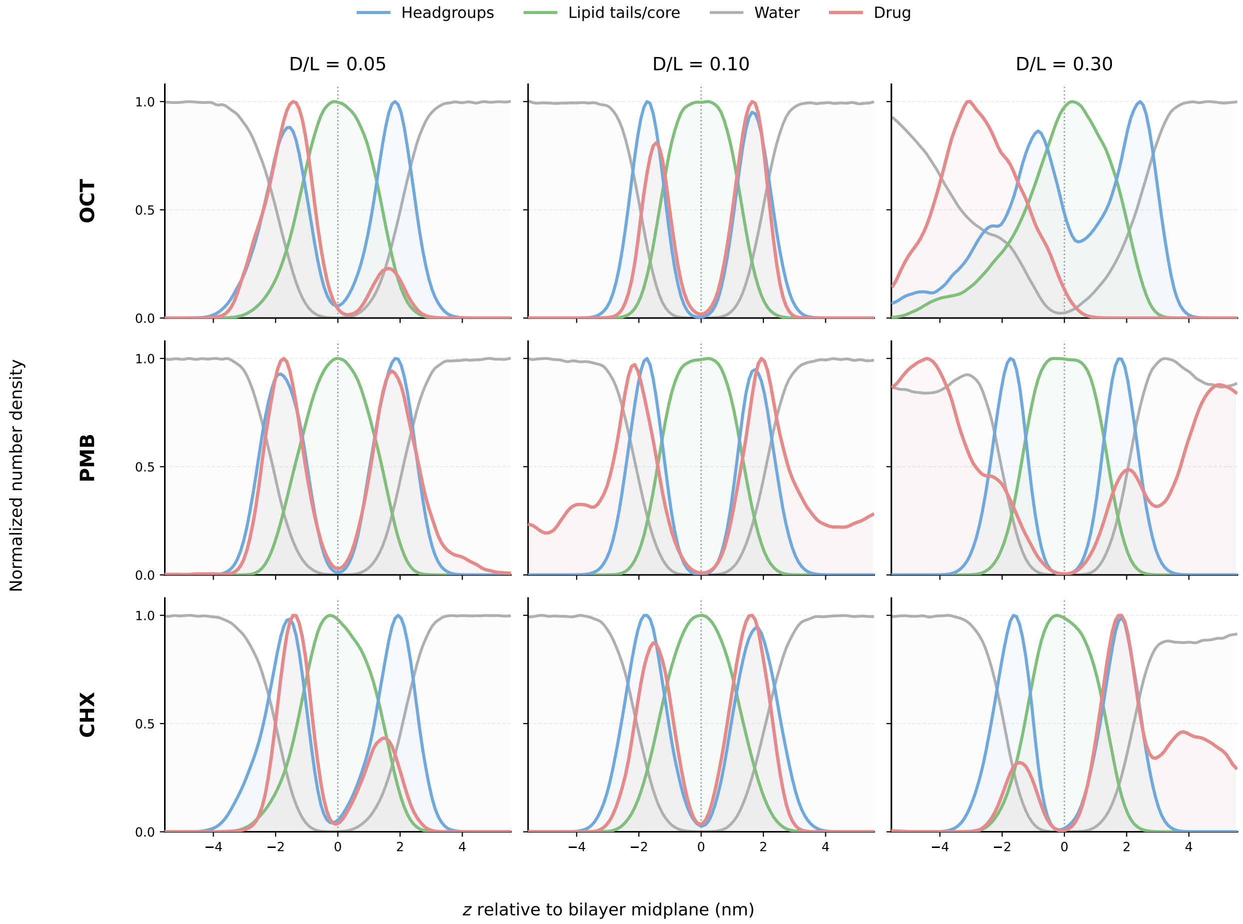


Figure 3: Normalized number-density profiles along the membrane normal for OCT (top row), PmB (middle row), and CHX (bottom row) at drug-to-lipid ratios of 0.05, 0.10, and 0.30 (left to right). Profiles are shown for lipid headgroups, the lipid-tail region, water, and antimicrobial molecules. The bilayer midplane is located at $z = 0$.

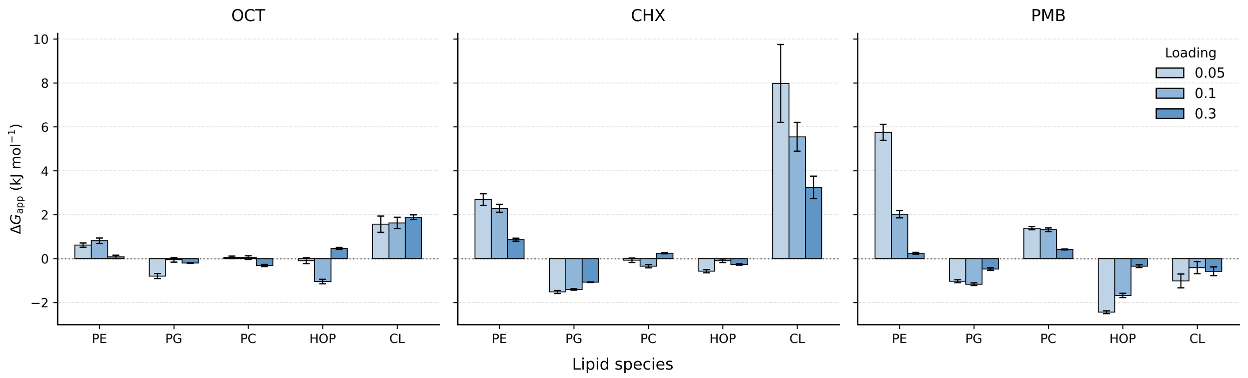


Figure 4: Lipid-resolved apparent compositional preference free energies, $\Delta G_{\mathrm{app},i}$, for (A) OCT, (B) CHX, and (C) PmB at drug-to-lipid ratios of 0.05, 0.10, and 0.30. Negative values indicate enrichment of a given lipid species within the local antimicrobial environment, whereas positive values indicate depletion. Error bars represent temporal uncertainties estimated using block averaging.

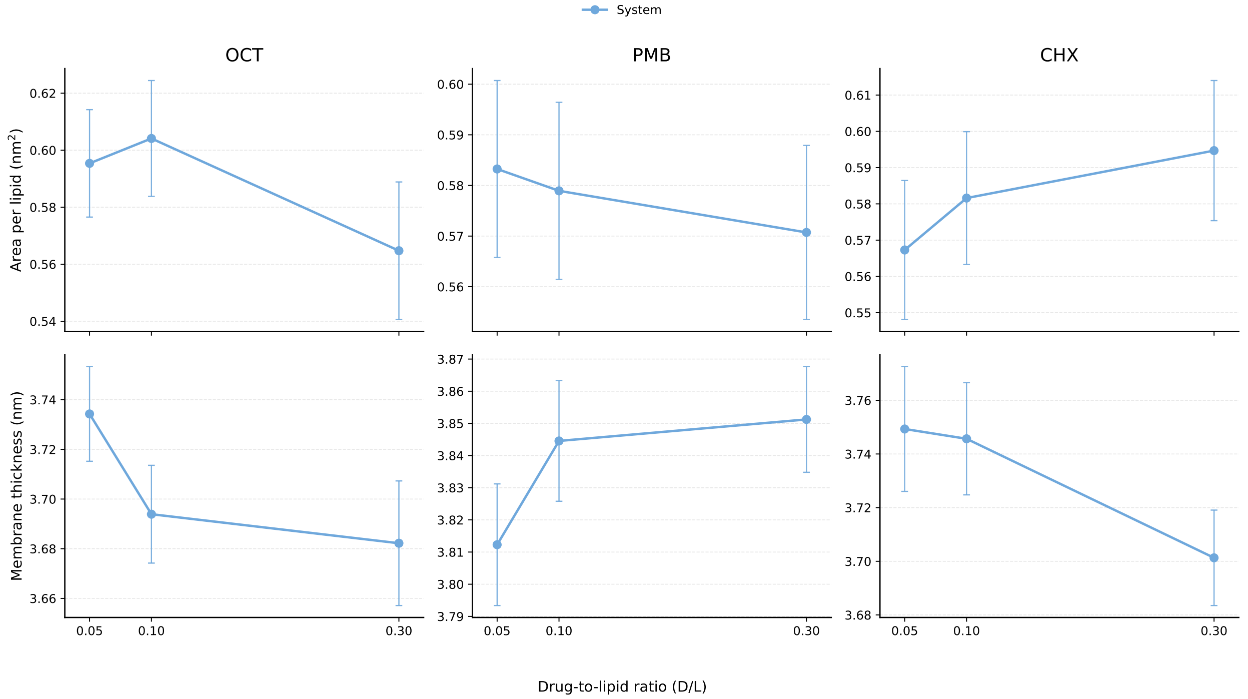


Figure 5: Global structural parameters of the simulated membranes. Area per lipid (top row) and membrane thickness (bottom row) are shown for OCT, PmB, and CHX at drug-to-lipid ratios of 0.05, 0.10, and 0.30. Points represent mean values over the final 500 ns of each trajectory, and error bars indicate temporal variability over the analysed interval.

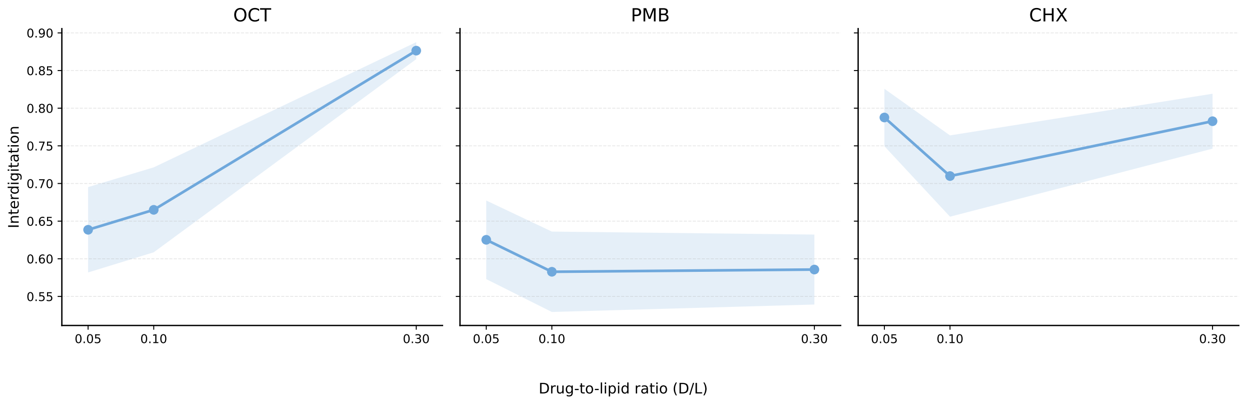


Figure 6: Interdigitation of the two membrane leaflets for OCT, PmB, and CHX at drug-to-lipid ratios of 0.05, 0.10, and 0.30. Symbols and connecting lines show mean values, while shaded regions indicate temporal variability over the final 500 ns of each trajectory.

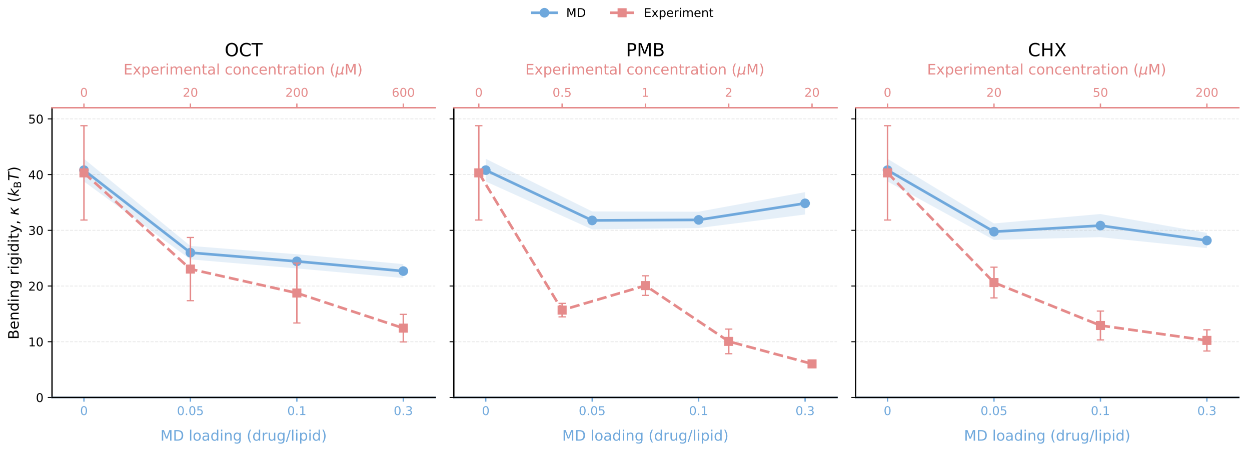


Figure 7: Absolute bending rigidity, $\kappa$, obtained from MD simulations (blue circles; lower x-axes) and GUV fluctuation spectroscopy (red squares; upper x-axes) for OCT (left), PmB (centre), and CHX (right). Simulation conditions are expressed as drug-to-lipid ratios, whereas experimental conditions are expressed as bulk antimicrobial concentrations.

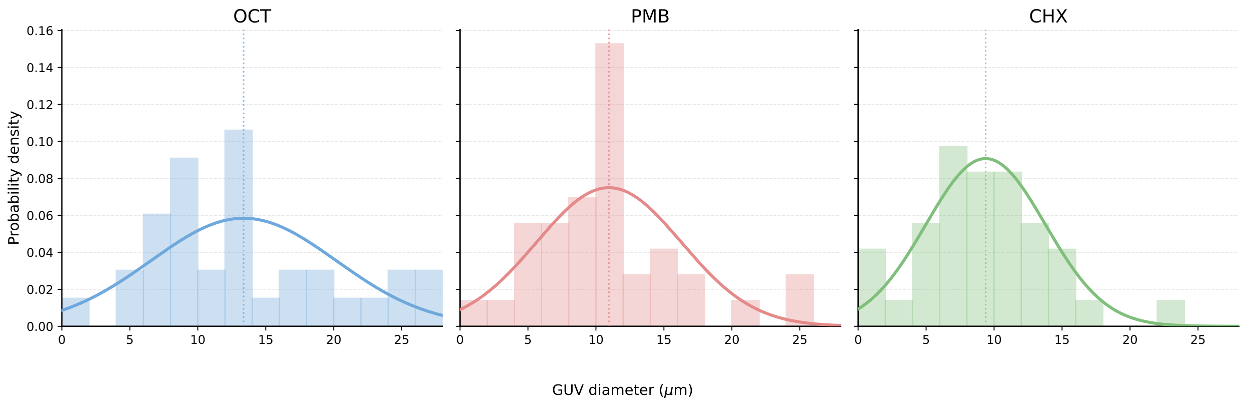


Figure 8: Diameter distributions of the GUVs analysed in the fluctuation-spectroscopy experiments with OCT (left), PmB (centre), and CHX (right). Bars show normalized diameter histograms, solid lines show kernel-density estimates, and vertical dotted lines indicate the mean diameter of each population.
